# Loss of function variants in DNAJB4 cause a myopathy with early respiratory failure

**DOI:** 10.1101/2022.07.31.502226

**Authors:** Conrad C. Weihl, Ana Töpf, Rocio Bengoechea, Jennifer Duff, Richard Charlton, Solange Kapetanovic Garcia, Cristina Domínguez-González, Abdulaziz Alsaman, Aurelio Hernández-Laín, Luis Varona Franco, Monica Elizabeth Ponce Sanchez, Sarah J Beecroft, Hayley Goulee, Jil Daw, Ankan Bhadra, Heather True, Michio Inoue, Andrew Findlay, Nigel Laing, Montse Olivé, Gianina Ravenscroft, Volker Straub

## Abstract

DNAJ/HSP40 co-chaperones are integral to the chaperone network, bind client proteins and recruit them to HSP70 for folding. We performed exome sequencing on patients with a presumed hereditary muscle disease and no genetic diagnosis. This identified four individuals from three unrelated families carrying an unreported homozygous stop gain (c.856A>T; p.Lys286Ter), or homozygous missense variants (c.74G>A; p.Arg25Gln and c.785T>C; p.Leu262Ser) in *DNAJB4.* Affected patients presented with axial rigidity and early respiratory failure requiring ventilator support between the 1^st^ and 4^th^ decade of life. Selective involvement of the semitendinosus and biceps femoris muscles was seen on MRI scans of the thigh. On biopsy, muscle was myopathic with angular fibers, protein inclusions and occasional rimmed vacuoles. DNAJB4 normally localizes to the Z-disc and was absent from muscle and fibroblasts of affected patients supporting a loss of function. Functional studies confirmed that the p.Lys286Ter and p.Leu262Ser mutant proteins are rapidly degraded in cells. In contrast, the p.Arg25Gln mutant protein is stable but failed to complement for DNAJB function in yeast, disaggregate client proteins or protect from heat shock induced cell death consistent with its loss of function. DNAJB4 knockout mice had muscle weakness and fiber atrophy with prominent diaphragm involvement and kyphosis. DNAJB4 knockout muscle and myotubes had myofibrillar disorganization and accumulated Z-disc proteins and protein chaperones. These data demonstrate a novel chaperonopathy associated with *DNAJB4* causing a myopathy with early respiratory failure. DNAJB4 loss of function variants may lead to the accumulation of DNAJB4 client proteins resulting in muscle dysfunction and degeneration.

## Introduction

Over 100 genes and proteins are associated with distinct neuromuscular phenotypes and pathologies. One group of neuromuscular disorders is hereditary myopathies. In the last decade the identification of novel genes associated with hereditary muscle diseases has rapidly expanded, driven by the application of whole exome sequencing (WES) to patients and their families. WES accompanied by deep-phenotyping and ancillary analysis of undiagnosed neuromuscular disease patients is critical to define new genetic myopathies^1^. For example, some genetically defined myopathies have specific associated symptoms such as facial weakness or respiratory involvement. Additionally, genetically similar myopathies may have distinctive myopathologic features such as protein inclusions or vacuoles observed in muscle biopsies. Finally, the use of muscle MRI can guide and support the diagnosis and assessment of genetically similar myopathies owing to selective muscle involvement that can be seen in many myopathies.

Myopathies associated with dominant or recessive mutations in protein chaperones have defined an emerging class of muscle diseases or chaperonopathies^2^. Protein chaperones facilitate the proper folding and degradation of client proteins within the cell. In the case of skeletal muscle, chaperones maintain sarcomeric structure during development and following rounds of repeated contraction^3^. Additionally, chaperones may recognize misfolded and aggregated proteins that are destined for degradation via autophagy or the ubiquitin proteasome system. Impaired chaperone function in skeletal muscle results in distinctive myopathology that includes myofiber atrophy, vacuolation, disordered internal architecture and protein inclusions^2^. One such disorder is associated with dominantly inherited mutations in the HSP40 cochaperone, DNAJB6, that cause limb-girdle muscular dystrophy type D1 (LGMD D1). DNAJB6 localizes to the Z-disc, and muscle tissue from patients with LGMD D1 has myofibrillar disorganization similar to that seen in patients with myofibrillar myopathy suggesting that DNAJB6 is important for sarcomere maintenance^4,5^

DNAJ/HSP40 co-chaperones are essential for chaperone-client specificity. They are grouped into three main categories DNAJA, DNAJB and DNAJC based on their domain similarities^6^. DNAJ proteins are highly conserved across species including lower eukaryotes. For example, the yeast DNAJB chaperone Sis1 is essential for yeast viability, and the human DNAJB1 can complement for its absence^7^. Dominant variants in *DNAJB6* and recessive mutations in *DNAJB2* cause a hereditary myopathy and neuropathy, respectively^4,5,8^. A single study of a multigenerational family has also been reported with dominant mutations in *DNAJB5* resulting in a clinical syndrome of distal atrophy^9^. DNAJB chaperones recognize misfolded client proteins and recruit HSP70 proteins. The DNAJB-client complex stimulates HSP70 ATP hydrolysis which then facilities the refolding of the client^6^. In the case of DNAJB6, it recognizes Z-disc proteins and recruits HSP70^10^. Dominant mutations in DNAJB6 trap HSP70 at the Z-disc resulting in an HSP70 dependent gain of function^10^. In the following study, we describe three families with homozygous loss of function variants in the HSP40 co-chaperone DNAJB4. Our data suggests that DNAJB4 similar to DNAJB6 is essential for myofibril maintenance and that aberrant chaperone function leads to isolated muscle disease.

## Methods

### Patient selection and evaluation

Patients from families A and B were recruited as part of the MYO-SEQ sequencing project^1^ and were sequenced using a 38-Mb targeted Illumina exome capture. Exome data was first screened for likely pathogenic variants using standard filtering criteria for rare diseases (MAF <1%, VEP moderate to high) and applying an *in silico* panel of 429 neuromuscular disease genes. DNA from the affected siblings in family C was exome sequenced using Ampliseq whole exome kit (ThermoFisher Scientific) and run on a Proton Sequencer, as described previously^11^. No relevant variants in genes known to be associated with neuromuscular conditions were found in any of the patients. Second round of analysis in order to identify novel candidate genes applied a more stringent criteria of high impact VEP and MAF<0.01%. *DNAJB4* variants were confirmed, and additional family members were segregated by Sanger sequencing.

### Muscle imaging

Muscle magnetic resonance images (MRI) and computer tomography (CT) scans were obtained on standard diagnostic scanners using T1-weighted axial scans, for patient A and B respectively.

### Yeast viability assay

Yeast strain used in this study was the derivative of Saccharomyces cerevisiae 74-D694 (ade1-14 his3-Δ200 leu2-3 112 trp1-289 ura 3-52). On this, Sis1 was deleted (sis1Δ::HygBMX4) as described previously1. Yeast cells were grown in rich media YPD (1% yeast extract, 2% peptone, 2% dextrose) or in synthetic defined (SD) media (0.67% yeast nitrogen base without amino acids, 2% dextrose) lacking specific nutrients to select for appropriate plasmids. Cells were transformed with plasmids pRS424-EV (empty vector), pRS424-HDJ1 (DNAJB1-WT), pRS424-HDJ1-R25Q (DNAJB1-R27Q), pRS414-Sis1WT, and pRS414-Sis1-R27Q using polyethylene-glycol/lithium-acetate (PEG/LioAC) technique2 and were selected using SD-trp plates. For yeast spotting’s, cultures were grown overnight in YPD. Cultures were pelleted, washed, and suspended in water to an optical density of 0.1. The normalized yeast solutions were pipetted into a 96-well plate, and serial dilutions (1:5) were made using a multichannel pipette. Finally, they were spotted onto plates containing YPD and also into medium containing 1mg/mL 5-fluoroorotic acid (5-FOA) that selects against cells maintaining URA3-marked plasmids and was used to replace WT Sis1 with the mutant constructs using an ethanol-sterilized 48-pin replicator. Plates were incubated for 5 days at 30°C and assessed for growth.

### Generation of DNAJB4 knockout (KO) C2C12 myoblasts

DNAJB4 KO C2C12 cells were generated using two guide RNAs (XCA1121b.m.Dnajb4.sp7 and XCA1121b.m.Dnajb4.sp8) based on off-target profile and the distance to the target site targeting two introns of *DNAJB4* to generate a 4-kb out of-frame deletion. Clones were screened for homozygosity of the 4-kb deletion via sequencing. Absence of DNAJB4 protein was confirmed via western blot.

### Antibodies

Antibodies used were the following: anti-rabbit GAPDH (Cell Signaling, 2118), anti-mouse Desmin (Dako, M0760), anti-rabbit DNAJB6 (Abcam, ab198995), anti-mouse HspA1 (Enzo, ADI-SPA-810), anti-rabbit α-actinin (Abcam, ab68167), anti-rabbit Myotilin (Abcam, ab78492), anti-rabbit Synemin (Bioss, bs-8555R), anti-rabbit αβ-crystallin (Enzo, ADI-SPA-223), anti-rabbit DNAJB4 (proteintech, 13064-1-AP) and anti-human DNAJB4 (Santa Cruz, sc-100711). Rhodamine Phalloidin Reagent (Invitrogen, PHDR1) was used to stain actin filaments in C2C12 myotubes for fluorescence microscopy. Secondary antibodies include anti-mouse HRP (Pierce), anti-rabbit HRP (Cell Signaling) anti-rabbit AlexaFluro (555 and 488).

### Plasmids and constructs

Mammalian constructs of *DNAJB4* were cloned using site-directed mutagenesis, digested with AflII/XhoI, and ligated into vector pcDNA3.1 containing a green fluorescent protein (GFP) tag. DNAJB4 R25Q, L262S, K286Ter mutations were generated with the Quick Change Mutagenesis Kit (Agilent Technologies #200517). Human TDP-43 fused to pCherry was described previously^10^.

### Immunoblotting

Muscle tissues and cultured cells were homogenized using RIPA lysis buffer (50 mM Tris–HCl, pH 7.4, 150 mM NaCl, 1% NP-40, 0.25% Na-deoxycholate and 1 mM EDTA) supplemented with protease inhibitor cocktail (Sigma-Aldrich), and lysates were centrifuged at 13 000 rpm for 10 min. Protein concentrations were determined using a BCA protein assay kit (Thermo Fisher Scientific). Aliquots of lysates were solubilized in Laemmli sample buffer and equal amounts of proteins were separated on 12% SDS-PAGE gels. Proteins were transferred to nitrocellulose membrane and then blocked with 5% nonfat dry milk in PBS with 0.1% Tween-20 for 1 h. The membrane was then incubated with primary antibodies, specific to the protein of interest, in 5% nonfat dry milk in PBS with 0.1% Tween overnight at 4°C. After incubation with the appropriate secondary antibody conjugated with horseradish peroxidase, enhanced chemiluminescence (GH Healthcare, UK) was used for protein detection. Immunoblots were obtained using the G:Box Chemi XT4, Genesys Version 1.1.2.0 (Syngene). Densitometry was measured with imageJ software (National Institute of Health).

### Solubility assay

C2C12 cells (differentiated for 5 days into myotubes), were collected and homogenized in 250 μl of 2% SDS-radioimmunoprecipitation assay buffer (50 mg Tris—Cl (pH 8.0), 150 mg NaCl, 1% NP-40, 0.5% sodium deoxycholate and 2% SDS) and protease inhibitor cocktail (Sigma-Aldrich). Homogenates were precleared with a 30s low speed spin and an aliquot of the supernatant was collected and named the total fraction. The additional supernatant was centrifuged at 100 000 × g for 30 min at 4°C and this supernatant was collected and named the soluble fraction. The pellet was then sonicated on ice after the addition of 150 μl of 5 M guanidine-HCl and re-centrifuged at 100 000 × g for 30 min at 4°C. This supernatant was removed and named the insoluble fraction. The insoluble fraction was precipitated by adding an equal volume of 20% trichloroacetic acid (Sigma-Aldrich) and incubated on ice for 20 min. Samples were then centrifuged at 10 000 × g for 15 min at 4°C and the resulting pellet was washed twice with ice-cold acetone. Residual acetone was removed by drying tubes at 95°C and the samples were resuspended in 50 μl of 5% SDS in 0.1N NaOH. The protein concentrations of all samples were determined using a BCA protein assay kit (Pierce). Each sample (30 μg) was analyzed by western blotting for each fraction.

### Animal generation and experimental protocols

*C57BL/6NJ-Dnajb4^em1(IMPC)J^*/Mmjax (DNAJB4 KO) strain was generated by the Knockout Mouse Phenotyping Program (KOMP) at The Jackson Laboratory using CRISPR technology. The targeted gene, DnaJ heat shock protein family (Hsp40) member B4 *(Dnajb4),* had an alteration that resulted in the deletion of 837 bp deletion in exon 5 and 274 bp of flanking intronic sequence, including the splice acceptor and donor, and is predicted to cause a change of amino acid sequence after residue 71 and early truncation 6 amino acids later. After the deletion of 381 bp of sequence there is a 6 bp [GGTCTT] endogenous retention followed by an additional 405 deleted bp. (Strain# MMRRC_046155-JAX). Animals were bred to C57/B6 (Jackson Laboratories) to at least the F5 generation. All animal experimental protocols were approved by the Animal Studies Committee of Washington University School of Medicine. Mice were housed in a temperature-controlled environment with 12 h light–dark cycles where they received food and water ad libitum. Mice were euthanized and skeletal muscle was dissected.

### Wire screen holding and grip test

Grip strength testing consisted of five separate measurements using a trapeze bar attached to a force transducer that recorded peak-generated force (Stoelting, WoodDale, IL, USA). Mice have the tendency to grab the bar with their forepaws and continue to hold while being pulled backwards by the tail, releasing only when unable to maintain grip. The resulting measurement was recorded and the average of the highest three measurements was determined to give the strength score. For every time point and strain, at least five animals were used. P-values were determined by a paired Student’s t-test. To validate our results, another quantitative strength measurement was performed by wire screen holding test. Mice were placed on a grid where it stood using all four limbs. Subsequently, the grid was turned upside down 15 cm above a cage. Latency for the mouse to release the mesh is recorded and the average hanging time of three trials was used as an outcome measure.

### In vivo electroporation

C57B6 (#:000664) mice (Jackson Labs, Bar Harbor, ME) were anesthetized using 2.5% isoflurane. We injected the TA (Tibialis anterior). The leg was saved and wiped with ethanol and then injected with endotoxin free plasmid diluted in sterile PBS to a volume of 50uL into FDB using a 29 gauge x 1/2 needle. Immediately after that, two-needle array electrodes were inserted longitudinally relative to myofibers. In vivo electroporation parameters for TA are as follows: voltage, 100 V; pulse length, 50 ms; number of pulses, six; pulse interval, 200 ms; desired field strength, 200 V/cm. Procedure was repeated on contralateral TA. Needles were removed and leg was wiped again with ethanol. Mice were then returned to recovery cage for monitoring.

### Cell culture, transfection, and heat shock

Flp-In T-REX 293 Cells (Invitrogen) expressing pcDNA5/FRT/TO constructs V5-DNAJB4_WT, V5-DNAJB4-R25Q, V5-DNAJB4-L262S, V5-DNAJB4-K286Ter were cultured in DMEM containing 4 mM l-glutamine (Invitrogen; 11965-084), 10% FBS (Atlanta Biologicals; S10350) and penicillin (50 IU)/streptomycin (50 μg/ml) (Invitrogen; 15 140), 50 μg/ml hygromycin B (Invitrogen; 10687-010), 50 μg/ml blasticidin (life technologies; R21001) and induced with 1 μg/ml tetracycline hydrochloride (Sigma; T76600) 48 h prior to experimentation. Cells were maintained in 5% CO2 at 37°C in 60 mm tissue culture-treated plates until the cells were 80— 85% confluent. HeLa cells were cultured in high glucose formulation of Dubelcco’s modified Eagle’s medium (Sigma-Aldrich) supplemented with 10% (vol/vol) fetal bovine serum, 2 mM L-glutamine and penicillin/streptomycin. C2C12 cells were maintained in proliferation media (DMEM with 20% FBS, 50 μg/mL P/S) and switched to differentiation media (DMEM with 2% horse serum, 50 μg/mL P/S) to form myotubes.

Transfection was performed with Lipofectamine 2000 (Life Technologies, Thermo Fisher Scientific; catalog 11668019) according to the manufacturer’s instructions. For heat shock experiment, HeLa cells were transfected either just with GFP-DNAJB4 plasmids or cotransfected with Ch-TDP43 constructs for 24 h. After transfection, cells were subjected to heat shock (42°C, 5% CO2) for 1 h.

### Histochemistry, immunohistochemistry and microscopy

Isolated muscle was mounted using tragacanth gum (Sigma, G1128) and quick frozen in liquid nitrogen-cooled 2-methylbutane. Samples were stored at −80°C until sectioning into 10-μm sections. Hematoxylin and eosin (H&E), nicotinamide adenine dinucleotide diaphorase (NADH), modified Gomori trichrome, myotilin, α-actinin and desmin staining was performed as previously described^5^. Sections were blocked in PNB (PerkinElmer), incubated with primary antibody followed by the appropriate secondary antibody. Briefly, muscle sections were affixed to slides incubated for 10 min in ice-cold acetone, mounted with Mowiol 4–88 (sigma) + DAPI and examined using a fluorescence microscope (Nikon 80i upright+ and Roper Scientific EZ monochrome CCD camera with deconvolution software analysis [NIS Elements, Nikon]). Non-fluorescent images were taken with a 5 megapixel color CCD (Nikon). Image processing and analysis were done with (NIS Elements 4.0) software and Adobe Photoshop. Fluorescent Images of random fields were taken with a ×20 objective using a Nikon Eclipse 80i fluorescence microscope. For electron microscopy, TA muscle was harvested, rinsed briefly in PBS and then placed immediately in Karnovsky fixative at 4°C for 24 h. Fixed muscle was embedded in plastic and sectioned for EM imaging with a JEOL JEM-1400 Plus 120 kV Transmission Electron Microscope equipped with an AMT XR111 high-speed 4 k × 2 k pixel phosphor-scintillated 12-bit charge coupled device camera. Image processing and analysis was done with (NIS Elements 4.0) software and Adobe Photoshop.

### X-ray

The X-ray images were taken with Faxitron^®^ Ultrafocus DXA system, a fully shielded x-ray cabinet designed for high-resolution imaging and dual-energy x-ray absorptiometry (DXA) analysis.

### Statistical analysis

Comparisons between two groups were made using a 2-tailed, paired Student’s t test. Comparisons between several groups were made using a 1-way ANOVA with Tukey’s post hoc test or Bonferroni’s correction to adjust for multiple comparisons. All analyses were performed with GraphPad Prism 8 (GraphPad Software). Data are presented as the mean ± SD, and results were considered statistically significant if P was less than 0.05. In some cases, the Student’s t test p value was calculated, and a Bonferroni-adjusted p value was determined for multiple comparisons.

## Results

### Homozygous variants in the HSP40 co-chaperone *DNAJB4* in patients with early respiratory failure

We report four affected individuals from three independent families clinically manifesting with respiratory failure associated with diaphragmatic weakness and spinal rigidity within the 1^st^ to 4^th^ decade of life. The clinical presentations are summarized in Table 1. *In silico* panel analysis did not identify any pathogenic changes in known neuromuscular disease genes. However, further analysis of the whole exome data identified a homozygous stop gain (c.856A>T; p.Lys286Ter) and two homozygous missense variants (c.74G>A; p.Arg25Gln and c.785T>C; p.Leu262Ser) in *DNAJB4* (NM_007034) in the affected individuals. Family members were tested and showed that the *DNAJB4* variants co-segregated with the disease and were inherited from each parent in Family A and C (parental DNA from Family B was not available) (Figure 1A). All heterozygous carriers of the *DNAJB4* variants were clinically unaffected. These variants are not reported in the Genome Aggregation Database (gnomAD; https://gnomad.broadinstitute.org/) and the affected missense residues are well conserved across species and had CADD scores of 28.8 and 33, respectively (Figure 1B).

**Figure 1:**
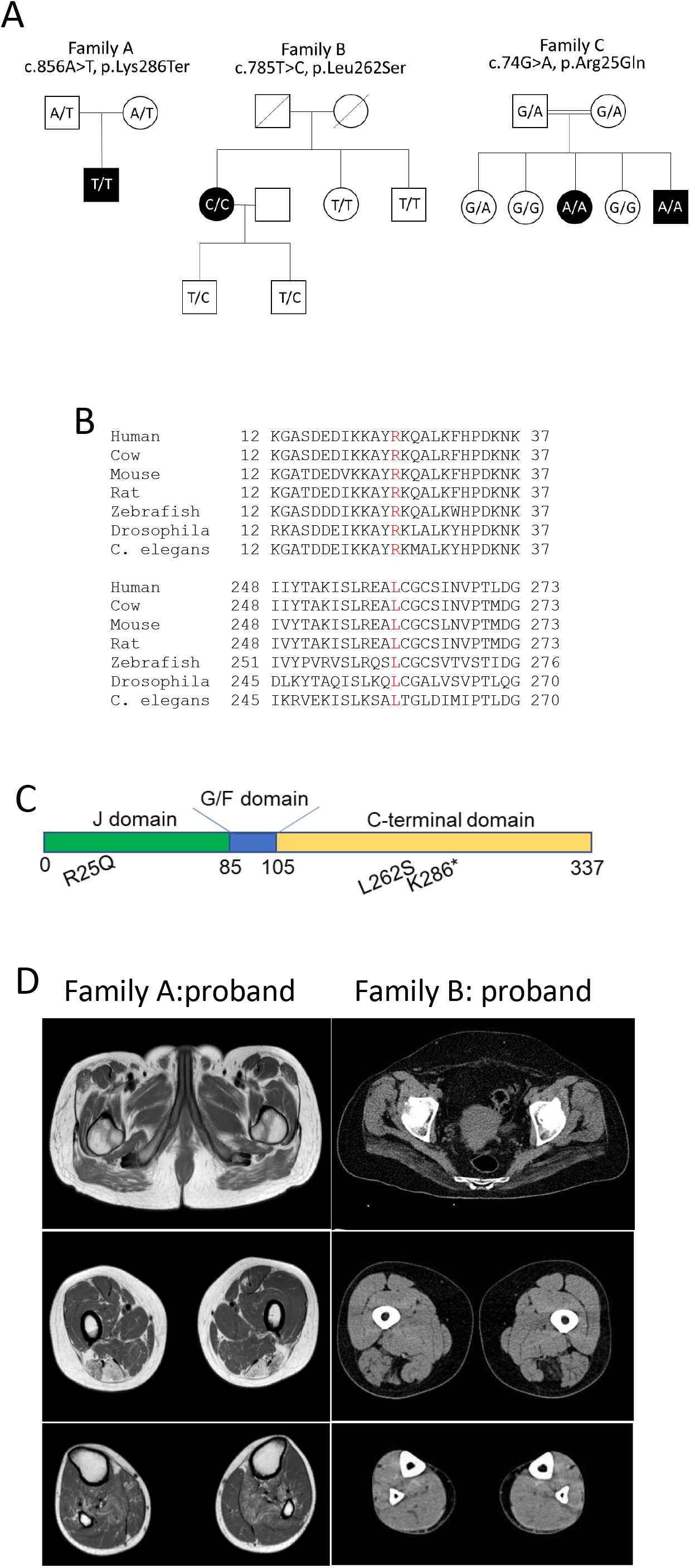
A) Pedigrees of three families with homozygous *DNAJB4* variants. Affected patients are in black. The genotype of the patient is represented within the pedigree. B) ClustalW alignment of DNAJB4 amino acid sequence from multiple species demonstrating conservation at R25 and L262. C) Schematic of the DNAJB4 denoting the J, G/F rich and C-terminal domains and the location of the identified variants. D) Lower extremity imaging of the probands from family A (MRI) and family B (CT scan).

The probands from Family A and Family B (PA:1 and PB:1) presented with respiratory failure in adulthood requiring non-invasive ventilator support. Patient PA:I was also noted to have spinal rigidity and axial weakness with a Beevor sign, and Patient PB:I was diagnosed with a hypertrophic cardiomyopathy. Two siblings from consanguineous parents, identified from Family C (PC:III and PC:V), presented with axial rigidity that included difficulty with bending and turning the neck at one year of age. Symptoms progressed to severe respiratory failure requiring invasive ventilation in patient PC:III and death in patient PC:V. There was no delay in motor milestones or evidence of limb weakness in any of the patients. Electromyography was consistent with myopathy and serum creatine kinase levels ranged from normal to 3X the upper limit of normal.

Imaging of the lower extremity showed a distinctive pattern of muscle disease in patients PA:I and PB:1 with selective involvement of the semitendinosus and biceps femoris muscles seen on MRI in PA:1 or by CT in PB:1 (Figure 1D). Imaging was not performed on patients from Family C. Muscle biopsies were obtained from patients PA:I and PB:I, and was noted to be myopathic with rare necrotic fibers (Figure 2A-B). Atrophic fibers were occasionally angular in isolation or grouped (Figure 2A-B). The biopsy from patient PA:I had additional pathological features including rimmed vacuoles and cytoplasmic inclusions seen on both hematoxylin and eosin and modified Gomori trichrome stains (Figure 2B-D). Many fiber regions had diminished oxidative enzyme activity and ATPase staining, especially in type 1 fibers (Figure 2E-F). These inclusions were also positive for desmin and myotilin by immunostaining (Figure 2G-H). Electron microscopy demonstrated Z-disc streaming and the accumulation of dense granulofilamentous material (Figure 2I-J).

**Figure 2:**
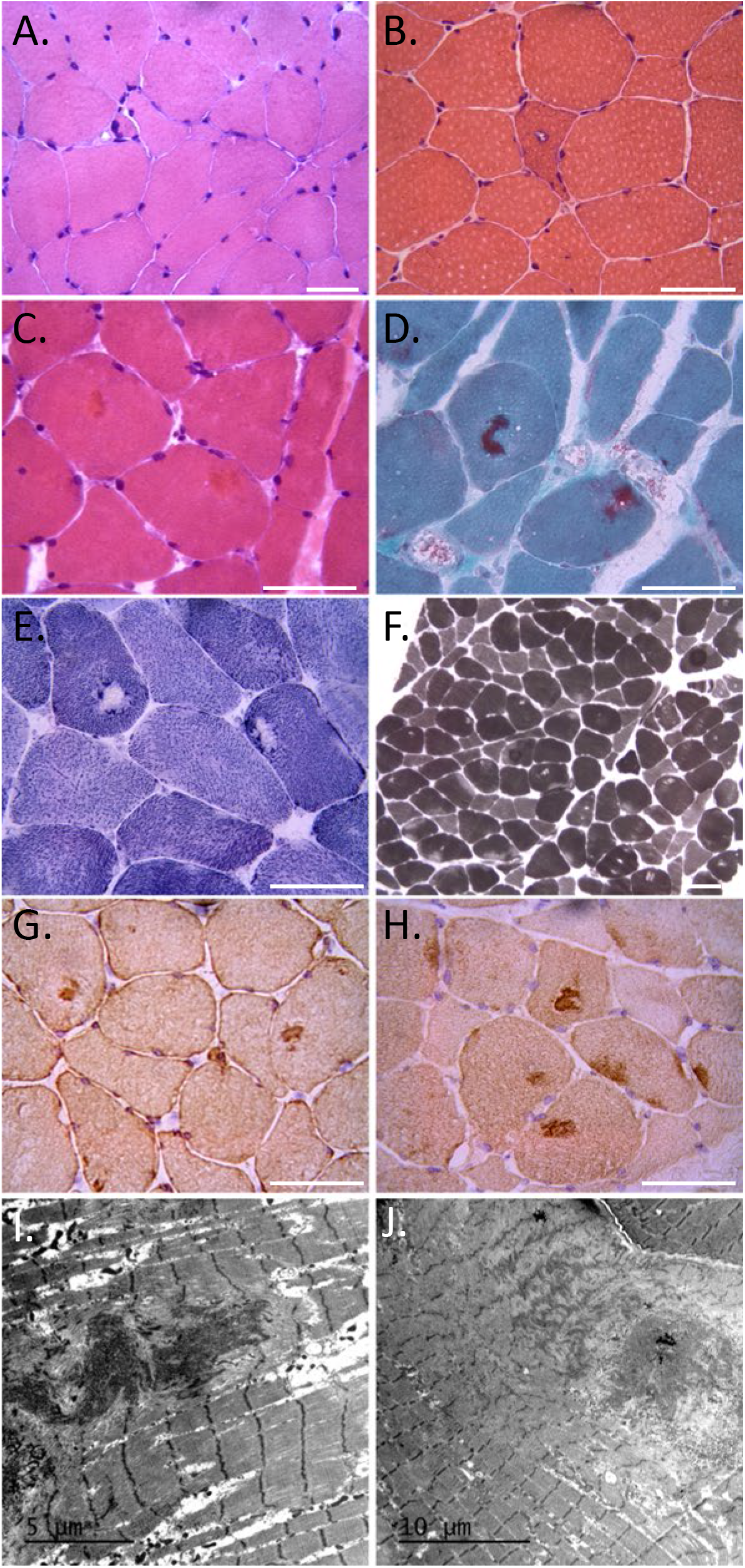
A) H&E staining of left biceps from patient B:I. B-C) H&E demonstrating a rimmed vacuole (B) and eosinophilic inclusions (C) from a deltoid biopsy of patient A:I. D) Modified Gomori trichrome staining of patient A:I demonstrating dark central inclusions. E) Many fiber regions have diminished oxidative enzyme activity (NADH) that sometimes is accentuated around the larger inclusions. F) These regions have also diminished had reduced ATPase staining, especially in type 1 fibers (ATPase 4,6). Scale bars are 50μM. G-H) Immunohistochemical staining for (G) desmin and (H) myotilin of muscle from patient A:I. I-J) Electron microscopy of muscle from patient A:I. Scale bars are 5 and 10μM respectively.

Since the c.856A>T; p.Lys286Ter nonsense variant, located in the last exon of *DNAJB4* is predicted to escape non-sense mediated decay, we performed an immunoblot for DNAJB4 using lysates from PA:I and control patient muscle. DNAJB4 migrated at its predicted molecular weight (~37kDa) in control muscle but was absent in PA:I (Figure 3A, B). A faint band was seen migrating between 25 and 37kDa suggesting that the truncated protein is translated but rapidly degraded (Figure 3C). Similarly, lysates from fibroblasts of PA:I and PB:I (c.785T>C; p.Leu262Ser) showed a decrease in total DNAJB4 protein levels suggesting that the p.Leu262Ser variant is also unstable (Figure 3B). We generated isogenic cell lines that stably express a tetracycline inducible V5 tagged DNAJB4 protein (V5-DNAJB4). Following washout of tetracycline, DNAJB4-WT is fully degraded by 3 days (Figure 3C). As expected, DNAJB4-K286Ter and DNAJB4-L262S were more rapidly degraded with the DNAJB4-K286Ter being truncated and both being absent by day 1 and 2 respectively (Figure 3C). In contrast, the DNAJB4-R25Q variant was more stable than DNAJB4-WT (Figure 3C). To see if the R25Q variant affected DNAJB4 localization in skeletal muscle, we performed electroporation of GFP-tagged DNAJB4-WT or DNAJB4-R25Q into the mouse footpad or mouse tibialis anterior (TA) muscle. In the footpad, DNAJB4-WT and DNAJB-R25Q localized to repeating structures that alternated with the A-band that was visualized by second harmonic generation imaging (Figure 3D). To confirm that this was the Z-disc, we immunostained similarly electroporated TA muscle with an antibody to α-actinin. DNAJB4-WT and DNAJB4-R25Q both co-localized to α-actinin supporting Z-disc localization of DNAJB4 (Figure 3E-F).

**Figure 3:**
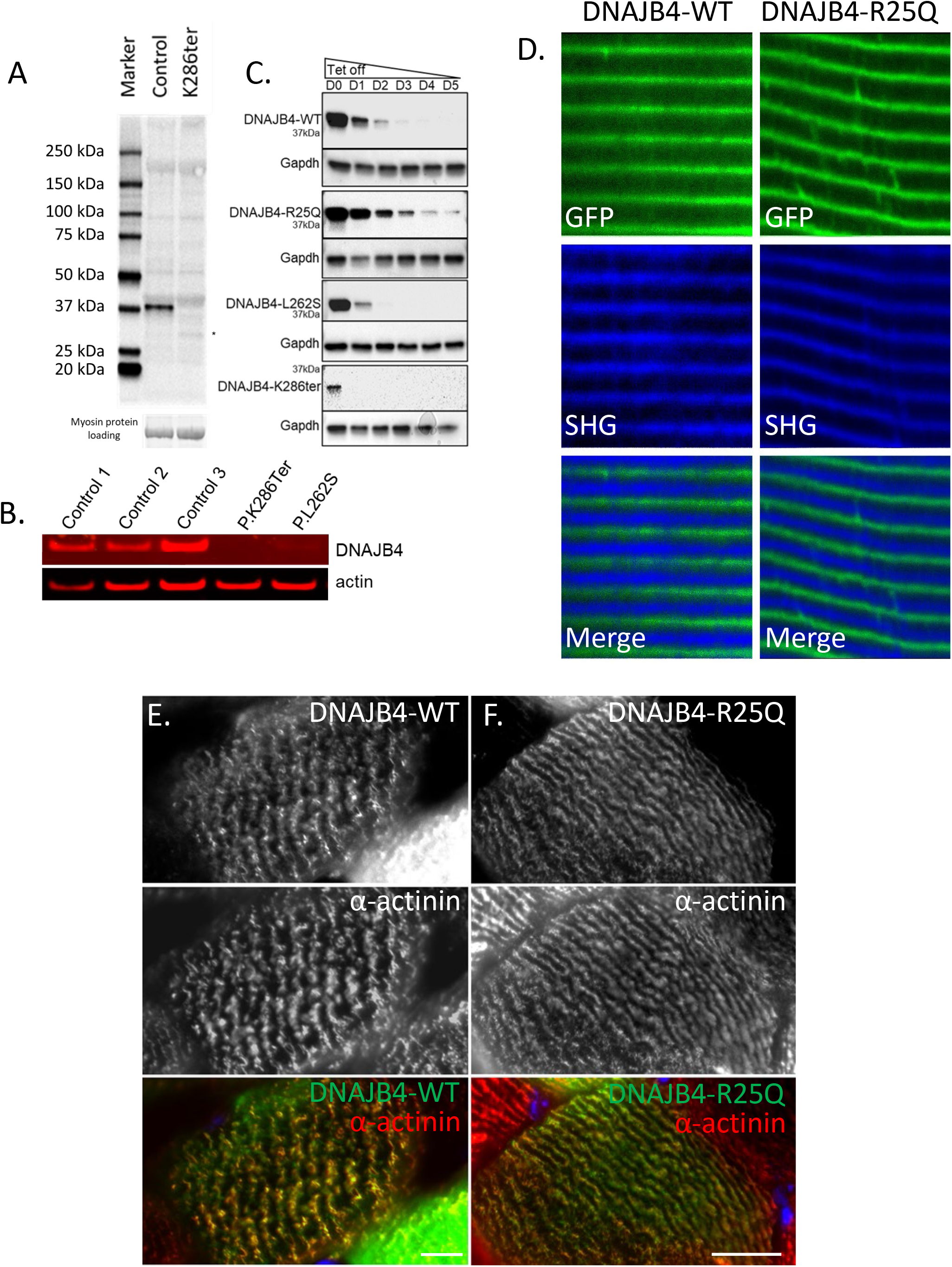
A) Immunoblot of skeletal muscle tissue lysates from control patient muscle or patient A:I carrying homozygous p.Lys286Ter variants in DNAJB4 with an anti-DNAJB4 antibody. B) Immunoblot from lysates of control patient fibroblasts or primary fibroblasts from patient A:I (p.Lys286Ter) and B:I (p.Leu262Ser) with anti-DNAJB4 or anti-actin antibodies. C) Isogenic 293T cells stably expressing V5 tagged DNAJB4-WT, DNAJB4-R25Q, DNAJB4-F90L, DNAJB4-L262S or DNAJB4-K286Ter under a tetracycline inducible promoter were treated with tetracycline for 48 hours and then lysates were harvested for five days following tetracycline removal. Lysates were then immunoblotted for V5 (DNAJB4) and GAPDH. D) Two-photon confocal microscopy of mouse footpad muscle electroporated with GFP tagged DNAJB4-WT or DNAJB4-R25Q (green) with associated second harmonic generation imaging (SHG). DNAJB4 is at the Z-disc as SHG marks the A-band. E-F) Immunofluorescent images of a single myofiber from mouse tibialis anterior muscle electroporated with a GFP tagged DNAJB4-WT (green) (D) or DNAJB4-R25Q (green) (E) and then immunostained with an antibody to α-actinin (red). Scale bar is 10μM.

To determine whether DNAJB4-R25Q had reduced function and would behave similar to the loss of function variants, DNAJB4-K286Ter and DNAJB4-L262S, we used a yeast complementation assay. All DNAJ proteins have a canonical J domain that defines their name. The J-domain is evolutionarily conserved and necessary for HSP70 interaction and ATPase activation^6^. Yeast have a homologous DNAJB protein, Sis1, essential for yeast viability and yeast prion propagation^7^. Mammalian DNAJB1 can complement for Sis1 function in yeast^7,10^ The R25 residue is well conserved across species and other DNAJB orthologues including Sis1 and DNAJB1 (Figure 4A). Notably, the R25 residue resides within helix II of the conserved J domain. Genetic replacement of yeast Sis1 with human DNAJB1-WT was sufficient to rescue yeast viability (Figure 4B). In contrast, replacement of Sis1 with DNAJB1-R25Q fails to complement, resulting in reduced or absent growth (Figure 4B). Similarly, generating the analogous R25Q mutation in yeast Sis1 (Sis1-R27Q) resulted in reduced viability as compared to Sis1-WT re-expression (Figure 4B). Human DNAJB4-WT fails to complement for loss of Sis1 in yeast (not shown).

**Figure 4:**
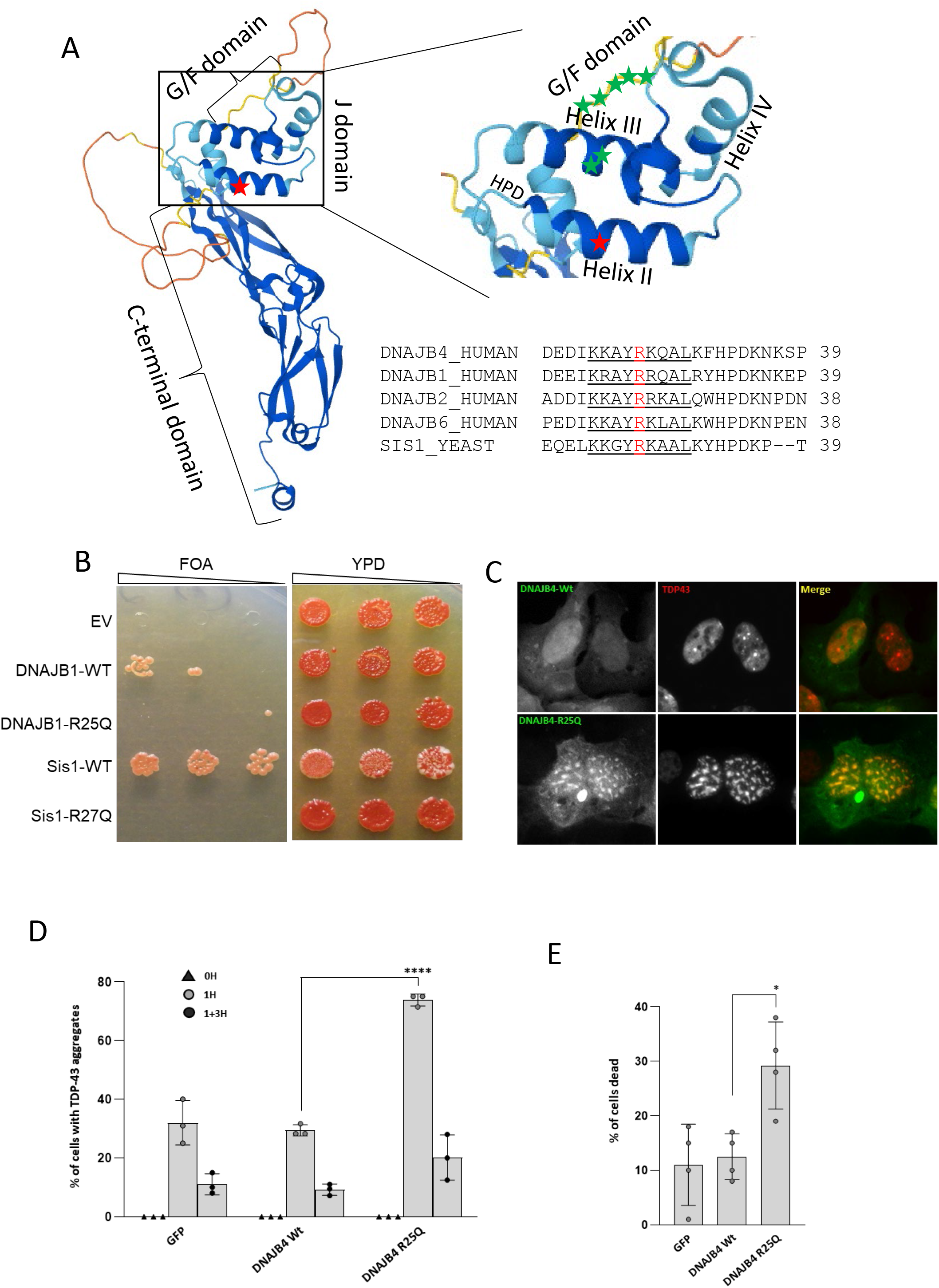
A) Rendering of DNAJB4 structure denoting the C-terminal, G/F and J domains with and enlargement of the J-domain showing Helix I-IV. Red star marks the R25 residue, green stars denote J domain residues (A50 and E54) and G/F domain residues (F89/F91/F93/N95/P96/D98/F100) mutated in DNAJB6 associated myopathy. Alignment of Helix II (underlined) from DNAJB4, DNAJB1, DNAJB2, DNAJB6 and Sis1. The R25 (DNAJB4/DNAJB1) and the R27 (Sis1) residue is in red. B) Yeast spottings from colonies that delete Sis1 and then complement with empty vector (EV), DNAJB-WT, DNAJB1-R25Q, Sis1-WT or Sis1-R27Q when plated on FOA media (left spottings). Spottings on full media (YPD) are on the right. C) Representative images of Hela cells transfected with GFP-DNAJB4-WT (green), or GFP-DNAJB4-R25Q (green) and mCherry-TDP-43 (red) one hour post heat shock. D) HeLa cells were co-transfected with GFP, GFP-DNAJB4-WT, or GFP-DNAJB4-R25Q and mCherry-TDP-43. The percentage of cells with TDP-43 nuclear inclusions at baseline (0H), post-heat shock (1H) or following heat shock recovery (1+3H) are represented graphically. E) HeLa cells were co-transfected with GFP, GFP-DNAJB4-WT, or GFP-DNAJB4-R25Q and mCherry-TDP-43 transfected with the indicated constructs, subjected to heat shock at 42°C for 1 hour, and the percentage of ethidium homodimer-1–positive cells was quantitated. Data are presented as the percentage of cells found positive/dead.

To explore the effect of DNAJB4-R25Q on DNAJB4 function in mammalian cells, we evaluated its ability to disaggregate a putative client, TDP-43. We have previously demonstrated that TDP-43 is a client protein for the DNAJB4 homolog DNAJB6 and that variants causing DNAJB6-associated myopathy disrupt this function^10^. Cells were co-transfected with mCherry-TDP-43 and GFP, GFP tagged DNAJB4-WT or GFP tagged DNAJB4-R25Q, and the percent of cells with TDP-43 nuclear stress granules were quantified at baseline, following a 1-hour heat shock at 42°C or 3 hours post heat shock recovery (Figure 4C-D). Notably, the DNAJB4-R25Q expressing cells had an increase in cells with persistent nuclear stress granules post-heat shock as compared with DNAJB4-WT expressing cells (Figure 4C-D). In a different assay, we evaluated cell death post heat shock on cells co-transfected with mCherry-TDP-43 and GFP, GFP tagged DNAJB4-WT or GFP tagged DNAJB4-R25Q (Figure 4E). Cells transfected with DNAJB4-R25Q had an increase in the percent of dead cells post heat shock consistent with a loss of function (Figure 4E).

The data above supported that the *DNAJB4* variants identified in our patients behaved as a loss of function. To understand whether a loss of DNAJB4 was detrimental to skeletal muscle, we generated and characterized DNAJB4 knockout mice. DNAJB4 homozygous knockout (DNAJB4-KO) mice were born at a normal Mendelian ratio, were viable and morphologically indistinguishable from control littermates. As expected, DNAJB4 was absent from all tested tissue lysates from DNAJB4-KO mice including TA, femoral quadriceps, gastrocnemius, and diaphragm muscles as compared with control mice (Figure 5A). DNAJB4-KO mice had kyphosis at 4 and 8 months of age (Figure 5B-C) and a decrease in the latency to fall on hanging grid testing after 4 and 8 months of age (Figure 5D). Notably, weakness was not appreciated using forelimb grip testing up to 8 months of age (Figure 5E). Hindlimb musculature that included the femoral quadriceps, TA and gastrocnemius muscles from DNAJB4-KO mice weighed less than wild-type controls at both 4 and 8 months (Figure 5F-G). Histopathology of TA, gastrocnemius and femoral quadriceps muscles of 4-and 8-month old DNAJB4-KO and wild-type controls revealed some myopathic features with scattered internal nuclei at 8 months and occasional myofibers with pale centers on NADH staining at 4 and 8 months (Figure 6A-B). Both muscle and myofiber atrophy were more apparent in DNAJB4-KO diaphragm muscle in which there was a reduction in diaphragm muscle thickness, myofiber atrophy and scattered fibers with core-like structures on NADH at 4 and 8 months (Figure 6C-D).

**Figure 5:**
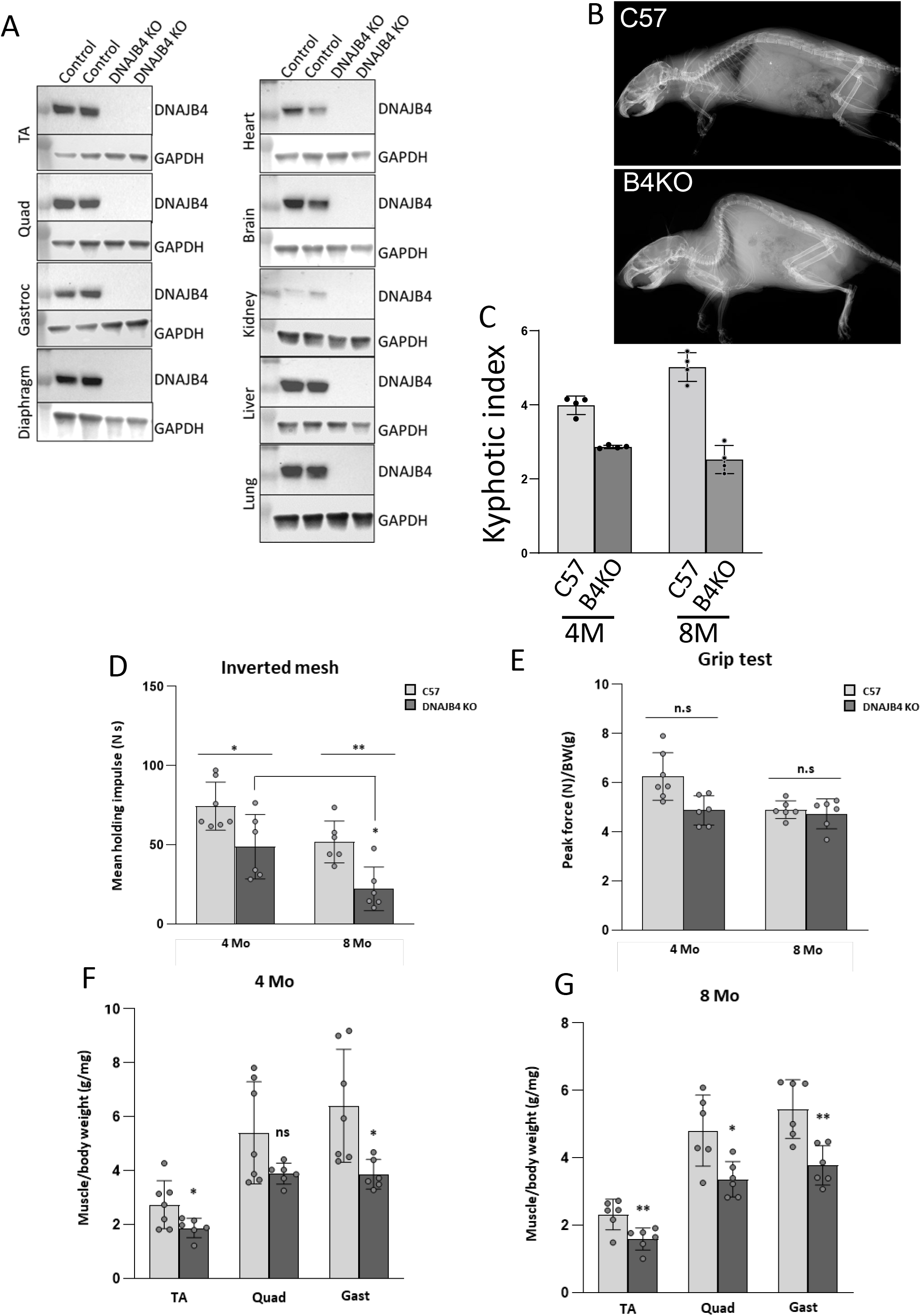
A) Lysates from skeletal muscle and tissues indicated from DNAJB4 homozygous knockout mice or control littermates immunoblotted with an antibody to DNAJB4 or GAPDH. B) X-ray image highlighting the skeleton of 8 month old DNAJB4 knockout (B4KO) or C57 control. C) Quantitation of the kyphotic index from 4 and 8 month old control or DNAJB4 knockout (B4KO) mice. D) Mean holding impulse on an inverted screen for 4 and 8 month old C57 control and DNAJB4 KO mice. E) Peak force forelimb grip strength testing for 4- and 8-month-old C57 control and DNAJB4 KO mice. F-G) Weight of indicated isolated muscles (tibialis anterior (TA), gastrocnemius (Gast) and quadriceps (Quad)) normalized to total body weight for 4- and 8-monthold C57 control and DNAJB4 KO mice.

**Figure 6:**
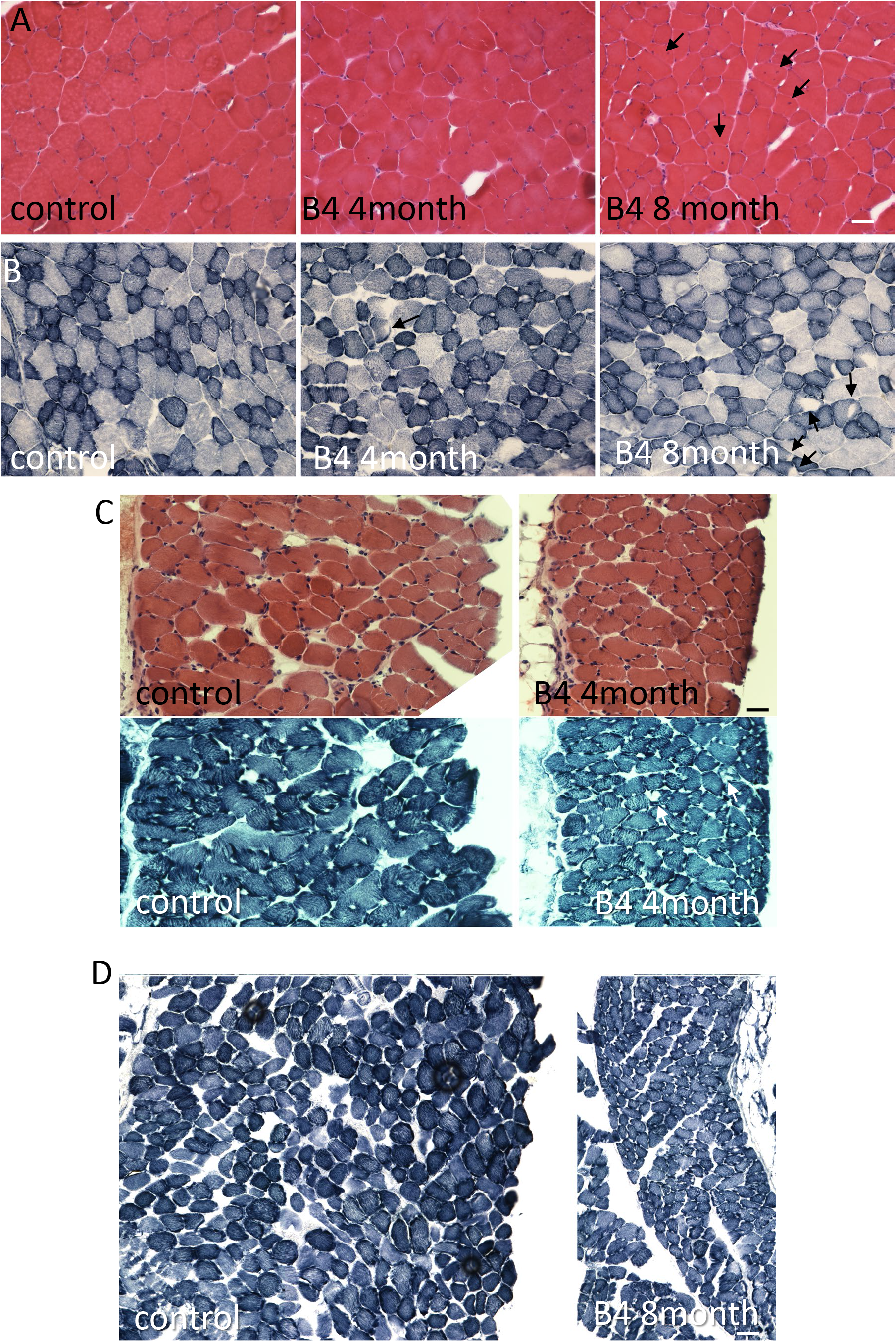
A) H&E staining of quadriceps muscle from 8-month-old control or 4- and 8-month old DNAJB4 KO mouse muscle. Fibers with internal nuclei are denoted with black arrows. B) NADH staining of tibialis anterior muscle from 8-month-old control or 4- and 8-month-old DNAJB4 KO mouse muscle. Fibers with central clearings are denoted with black arrows. C) H&E and NADH staining of diaphragm muscle from 4-month-old control or 4-month-old DNAJB4KO mice. Note myofiber atrophy and decreased diaphragm thickness. White arrows denote central clearings on NADH. D) NADH staining of diaphragm muscle from 8-month-old control or 8-month-old DNAJB4KO mice. Note myofiber atrophy and decreased diaphragm thickness. Scale bars are 50μM.

To further understand the role of DNAJB4 in skeletal muscle, we immunoblotted TA muscle lysates from 4-month old control and DNAJB4-KO mice. Remarkably, DNAJB4-KO muscle had an increase in the Z-disc proteins desmin and myotilin. Expression of α-actinin was unchanged, whereas the level of synemin was decreased (Figure 7A-B). The chaperone proteins HSPA1 and CRYAB were elevated whereas DNAJB6 levels were unchanged (Figure 7A-B). An increase in Z-disc proteins desmin, myotilin and α-actinin was seen in TA muscle lysates from 8-month old DNAJB4-KO mice as compared to control mice (Figure 7C-D).

**Figure 7:**
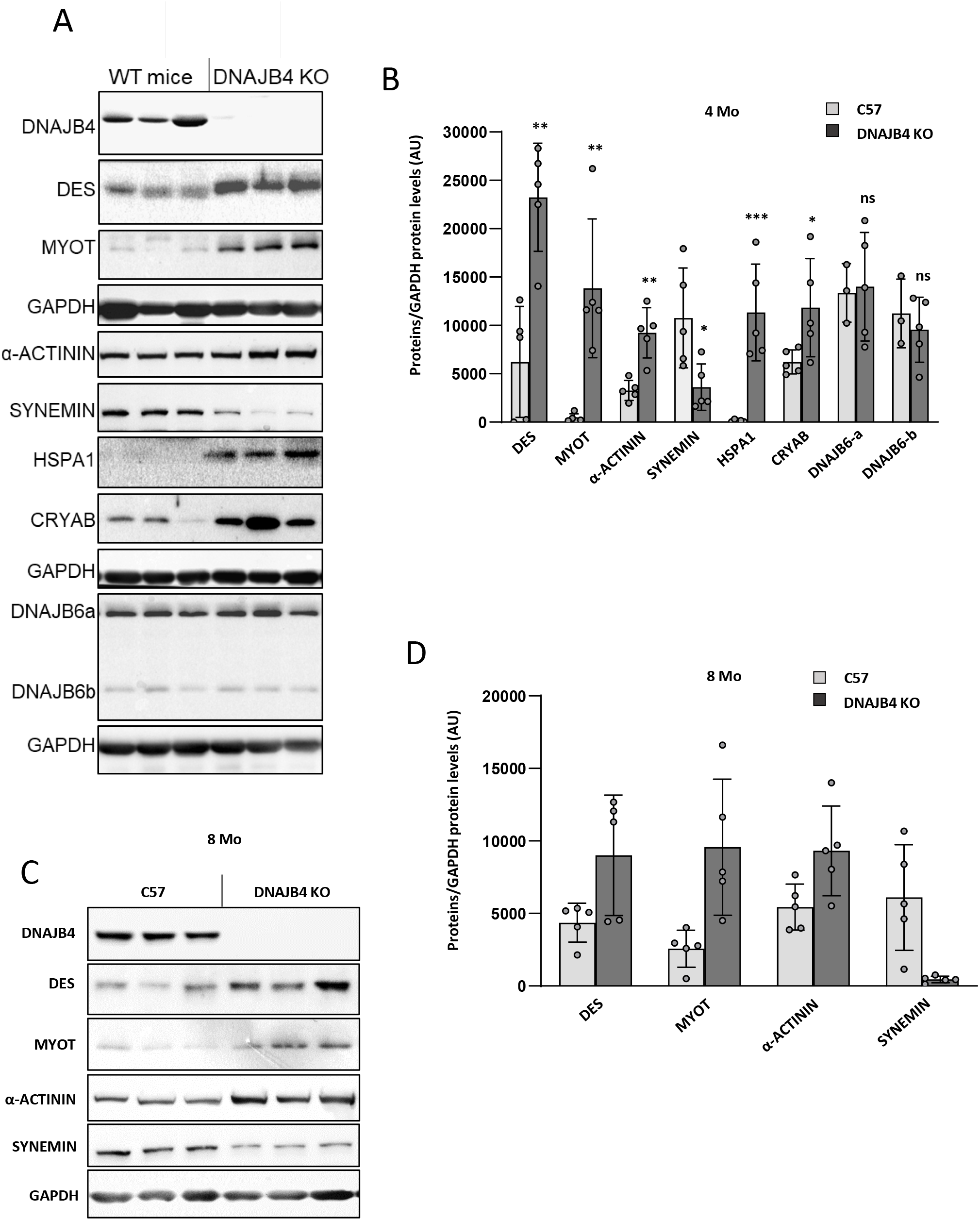
A) Immunoblots of lysates from 4-month-old mouse tibialis anterior of control or DNAJB4 knockout mice with antibodies to DNAJB4, desmin (DES), myotilin (MYOT), α-actinin, synemin, HSPA1, CRYAB, DNAJB6 or GAPDH. B) Quantitation of immunoblotted proteins from TA muscle at 4 months. N=5/condition. *, **, *** indicate p values 0.01, 0.001 and 0.0001. C) Immunoblots of lysates from 8-month-old mouse tibialis anterior of control or DNAJB4 knockout mice with antibodies to DNAJB4, desmin (DES), myotilin (MYOT), α-actinin, synemin or GAPDH. D) Quantitation of immunoblotted proteins from TA muscle at 8 months. N=5/condition. *, **, *** indicate p values 0.01, 0.001 and 0.0001.

We generated two independent C2C12-DNAJB4 KO (B4KO) cell lines using CRISPR/Cas9. B4KO myoblasts were indistinguishable from the parent C2C12 cells with regard to morphology and cell viability. Although the differentiation index as defined by nuclear incorporation into a myotube was similar, the number of myonuclei per myotube and the size of myotubes were greater consistent with increased fusion (Figure 8A-C). Immunoblotting of cell lysates from C2C12 myoblasts or myotubes following 5- and 10-days differentiation demonstrated an increase in desmin and α-actinin in B4KO cells as compared with the parental C2C12 myoblast line (Figure 8D). Ultracentrifugation of 5-day old myotube lysates to separate soluble and insoluble fractions demonstrated the presence of DNAJB6, CRYAB and desmin in the insoluble fractions of B4KO myotubes (Figure 8E). Immunofluorescence for myofibrillar proteins demonstrated aggregation of desmin, actin and myotilin that was increased in B4KO myotubes (Figure 8F-G).

**Figure 8:**
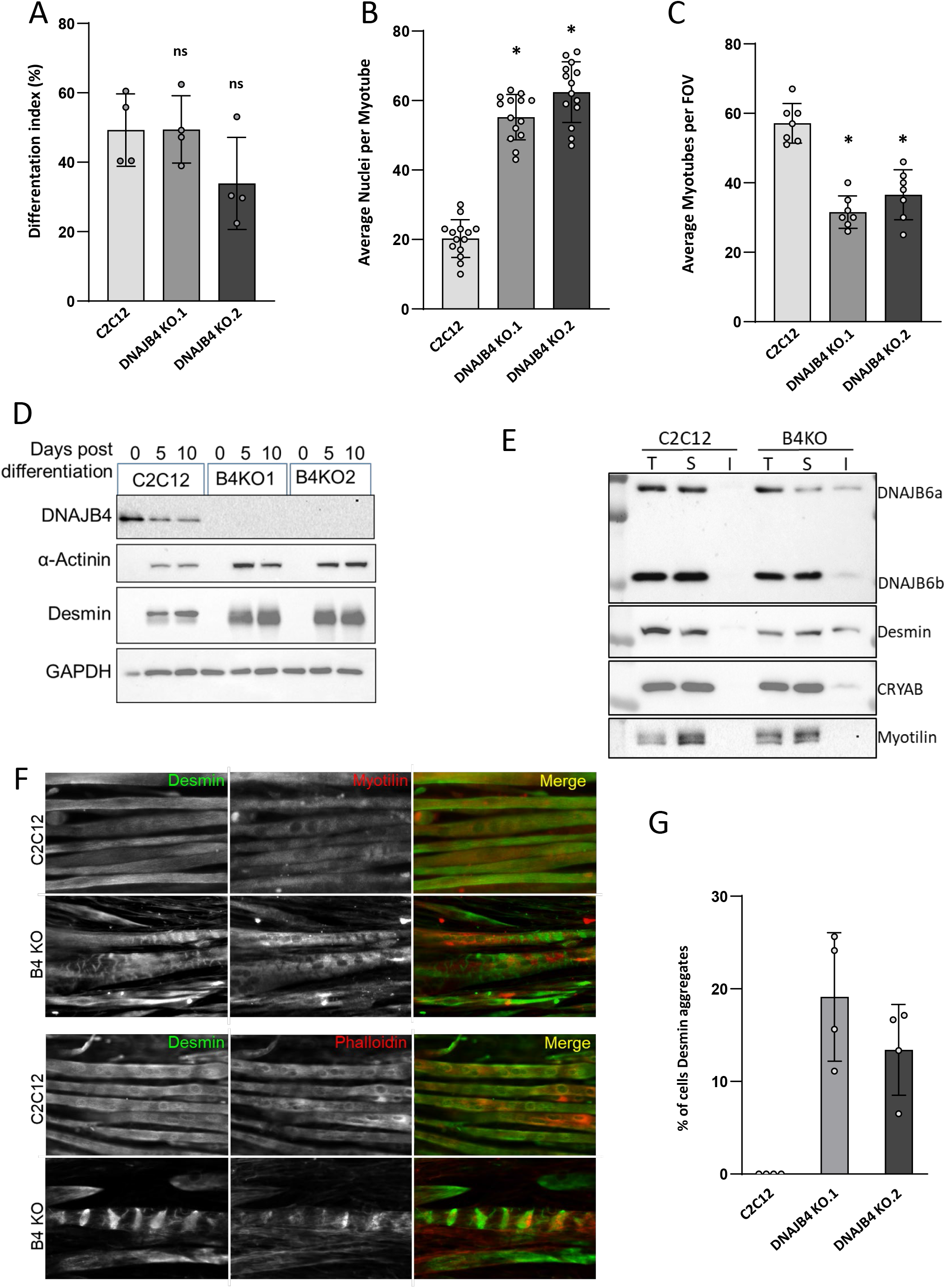
A-C) Control C2C12 or one of two different DNAJB4 knockout (B4KO) lines were placed in differentiation media for 5 days and then the differentiation index, number of nuclei/myotube and total number of myotubes was quantitated. D) Immunoblots of lysates from undifferentiated (0) or 5- and 10-days post differentiation of control or DNAJB4 knockout C2C12 cells (B4KO1 or B4KO2) with antibodies to DNAJB4, desmin, α-actinin or GAPDH. E) Immunoblots of lysates from differentiated (5 days) C2C12 or B4KO myotubes following detergent lysis and ultracentrifugation into a total (T) lysate, soluble (S) fraction and insoluble (I) fraction with antibodies to DNAJB6, desmin, CRYAB and myotilin. F) Immunofluorescent images of 5 day differentiated C2C12 or B4 KO myotubes using an antibody to desmin (green) and myotilin (red upper panels) or phalloidin (red lower panels). G) Graph of the percentage of fibers with desmin inclusions in control or DNAJB4 KO myotubes.

## Discussion

Our studies identify a new muscle disease associated with recessive variants in *DNAJB4.* Several lines of evidence support our assertion that pathogenic variants in the *DNAJB4* gene are causative in our patients. Specifically, the phenotypic similarities of early respiratory failure in all patients support a distinct clinical syndrome. The similar pattern of weakness is further substantiated by muscle imaging studies that reveal exclusive involvement of the semitendinosus and biceps femoris muscles. Finally, myopathology with protein inclusions and vacuoles similar to that seen in other myopathies associated with protein chaperone dysfunction function supports a defect in *DNAJB4* activity.

Other myopathies associated with early and sometimes isolated respiratory failure include late-onset Pompe disease, multi-minicore myopathy due to *SEPN1* variants, myofibrillar myopathies, hereditary myopathy with early respiratory failure (HMERF) associated with dominant mutations in *TTN*^12^ and nemaline myopathies caused by *TNNT1* variants^13^. These hereditary myopathies all have some pathologic features similar to *DNAJB4* patients with myofibrillar disorganization, protein inclusions and autophagic vacuoles suggesting that respiratory muscles such as the diaphragm are more sensitive to proteostatic stress^12^.

The *DNAJB4* patients described in this report resembled previous descriptions of HMERF patients associated with *TTN* variants, those with *TNNT1* variants and some patients with *DES* and *CRYAB* mutations. Specifically, these patients have respiratory failure as the presenting symptom without obligatory associated cardiomyopathy^14–16^. In addition, and as described for *DNAJB4, TNNT1* patients may also present with rigid spine. On muscle MRI, both HMERF and *DNAJB4* myopathies have early and unappreciated atrophy of the semitendinosus a feature also observed in *DES* or *CRYAB* patients^17^. However, distinct from HMERF which spares the adjacent biceps femoris, this muscle is also affected in our *DNAJB4* patients^15^. As mentioned, the myopathology in *DNAJB4* patients has eosinophilic inclusions that are desmin and myotilin positive with other myofibers containing rimmed vacuoles. These distinctive pathologies are similarly seen in HMERF, and *TNNT1* patients also show areas containing eosinophilic inclusions, and nemaline rods that are myotilin positive. However, under electron microscopy, in *DNAJB4* patients, prominent sarcomeric disorganization, core formations, and accumulation of granulofilamentous material intermingled with dense electron dense filaments were observed. It is important to note that HMERF patients typically show collections of cytoplasmic bodies seen as eosinophilic inclusions that stain darkly on Gomori trichrome and are often located at the at the periphery of muscle fibers^15^. Our patients did not have any appreciable cytoplasmic bodies or nemaline rods on their biopsies.

Dominant mutations in other chaperones such as *DNAJB6* or *BAG3* cause hereditary myopathies with biopsies resembling myofibrillar myopathy^4,5,18^. In these myopathies, a gain of function or dominant interaction with wild-type chaperones is thought to be the pathogenic mechanism^10,19^. Similarly dominant mutations in the small HSP, *CRYAB* result in a late onset myofibrillar myopathy with cardiac dysfunction and cataracts^20^. On the other hand, homozygous loss of function mutations in *CRYAB* are associated with a severe infantile onset myopathy with respiratory insufficiency and myofibrillar changes on biopsy^16^. The discrepancy in disease severity between dominant and recessive mutations may be explained by residual chaperone activity since dominant patients retain WT protein.

Our patients carried three different homozygous variants in *DNAJB4.* Two patients were homozygous for the missense variant (p.Leu262Ser) or the stop-gain variant (p.Lys286Ter) and had adult onset respiratory failure. Functional studies demonstrated that these variants destabilized the protein resulting in its rapid degradation and absence from patient skeletal muscle and fibroblasts. Two siblings with childhood onset respiratory failure were homozygous for a *DNAJB4* missense variant within helix II of the J domain (p.Arg25Gln). Unlike the L262S and K286Ter variants, DNAJB4-R25Q was stable in cells, yet yeast complementation studies confirmed that the DNAJB4-R25Q variant was non-functional. Yeast complementation assays with the homologous dominant mutations in DNAJB6 residing in the G/F domain or helix III of the J domain also failed to rescue viability^10,21^. In the case of expression of DNAJB6 G/F-domain mutations, viability was rescued by abrogating its interaction with HSP70, suggesting a dominant effect on HSP70 function^10^. Whether the DNAJB4-R25Q variant behaves as a loss of DNAJB4 function, a toxic gain function through interactions with HSP70 or a combination of both remains to be determined. However, it is intriguing to speculate that the more severe phenotype is related to both a loss of function and gain of toxicity by DNAJB4.

DNAJB4 is ubiquitously expressed yet loss of function leads to a myopathy in patients and knockout mice. Skeletal muscle may be particularly vulnerable to chaperone dysfunction as mutations in many ubiquitously expressed chaperones lead to myopathy. For example, contractile stress on the myofibrillar apparatus may lead to protein unfolding and aggregation that eventually overwhelms the proteostatic network^3^. It is well established that IgG like repeats in titin fold and unfold during muscle contraction and that protein chaperones facilitate this process^22^. It is also possible that DNAJB4 chaperones a specific set of client proteins that are essential for muscle function. For example, a BAG3/HSPB8 chaperone complex recognizes unfolded filamin C triaging it for degradation^23^. Similarly, the myosin specific chaperone unc-45 along with HSP70 and HSP90 facilitate the assembly of myosin filaments and the periodicity of myosin heads^24^. Notably mutations in both *BAG3* and *UNC45* lead to myopathies with disorganized myofibrillar structures^18,25^. Whether DNAJB4 has a select set of clients remains to be established.

## Funding

The results reported here were generated using funding received from the Solve-RD project within the European Rare Disease Models & Mechanisms Network (RDMM-Europe). The Solve-RD project has received funding from the European Union’s Horizon 2020 research and innovation programme under grant agreement No 779257. MYO-SEQ was funded by Sanofi Genzyme, Ultragenyx, LGMD2I Research Fund, Samantha J. Brazzo Foundation, LGMD2D Foundation and Kurt+Peter Foundation, Muscular Dystrophy UK, and Coalition to Cure Calpain 3. Analysis was provided by the Broad Institute of MIT and Harvard Center for Mendelian Genomics (Broad CMG) and was funded by the National Human Genome Research Institute, the National Eye Institute, and the National Heart, Lung, and Blood Institute grant UM1 HG008900, and in part by National Human Genome Research Institute grant R01 HG009141. CCW is funded by R01AR068797 and K24AR073317. This research was also supported by a grant from the Australian NHMRC (APP2002640). GR is supported by an Emerging Leader Fellowship from the NHMRC (APP2007769). MO is supported by a grant from the Spanish Ministry of Health, Fondos FEDER-ISCIII PI21/01621. CD, AHL and MO are members of the ERN EURO-NMD.

## Notes

### Competing Interest Statement

The authors have declared no competing interest.

### Summary of Updates

Authorship changed

